# Can the creation of new freshwater habitat demographically offset losses of Pacific salmon from chronic anthropogenic mortality?

**DOI:** 10.1101/2020.07.21.213553

**Authors:** Pascale Gibeau, Michael J. Bradford, Wendy J. Palen

## Abstract

Over 1 billion USD are devoted annually to rehabilitating freshwater habitats to improve survival for the recovery of endangered salmon populations. Mitigation often requires the creation of new habitat (e.g. habitat compensation) to offset population losses from human activities, however compensation schemes are rarely evaluated. Anadromous Pacific salmon are ecologically, culturally, and economically important in the US and Canada, and face numerous threats from climate change, over-harvesting, and degradation of freshwater habitats. Here we used a matrix population model of coho salmon *(Oncorhynchus kisutch)* to determine the amount of habitat compensation needed to offset mortality (2-20% per year) caused by a range of development activities. We simulated chronic mortality to three different life stages (egg, parr, smolt/adult), individually and in combination, to mimic impacts from development, and evaluated if the number of smolts produced from constructed side-channels demographically offset losses. We show that under ideal conditions, the typical size of a constructed side-channel in the Pacific Northwest (PNW) (3405 m^2^) is sufficient to compensate for only relatively low levels of chronic mortality to either the parr or smolt/adult stages (2-7% per year), but populations do not recover if mortality is >10% per year. When we assumed lower productivity (e.g.; 25^th^ percentile), or imposed mortality at multiple life stages, we found that constructed channels would need to be larger (0.2-4.5 times) than if we assumed mean productivity or as compared to the typical size built in the PNW, respectively, to maintain population sizes.. We conclude that habitat compensation has the potential to mitigate chronic mortality to early life stages, but that current practices are likely not sufficient when we incorporate more realistic assumptions about productivity of constructed side-channels and cumulative effects of anthropogenic disturbances on multiple life stages.

## Introduction

Billions of dollars are spent annually to mitigate impacts of human activities on biotic communities and abiotic processes. The stakes of such ecological and economic trade-offs are high, as total investments in energy, water, and infrastructure development projects are expected to exceed $53 trillion (US) worldwide between 2010 and 2030 (OECD 2012 in Tallis et al. 2015). Regulatory agencies often require that developers apply the mitigation hierarchy, which consists of avoidance, then minimization, and finally compensation or offset for the impact of projects on ecosystems and biodiversity (Quétier and Lavorel 2011; Tallis et al. 2015). In many jurisdictions, compensation activities are often guided by a requirement for “No Net Loss” of biodiversity, where losses from development are required to be fully offset such that population sizes are maintained and stable after development (Bull et al. 2013). To guide biodiversity compensation, managers develop measures, called equivalency metrics, which estimate the amount of compensation needed to offset losses due to development. In practice, the difficulty of developing metrics that address impacts to biodiversity broadly (Bull et al. 2013) results in compensation targeting a single, economically important, species, or is measured in terms of lost habitat, individuals, or productivity of populations. Equivalency metrics are particularly challenging to develop given that losses from development actions are difficult to predict in advance, while gains from compensation are often delayed, leading to uncertainty in the outcomes (Bradford 2017). Additionally, compensation activities have been criticized as doing more harm than good by creating the perception that negative consequences from development can be effectively mitigated despite few successful examples (Moore and Moore 2013).

Despite their shortcomings, compensation activities and offsetting are mandated through legislation in 45 countries, and are under development in another 27 (Bull et al. 2013). For example, the US *Clean Water Act* Section 404 (1972) includes a program that requires mitigation for all activities that impact fish or wildlife habitat and the use of the mitigation hierarchy (Doyle and Shields 2012). Similarly, in Canada the *Fisheries Act* includes a provision to employ offsetting since 2015 as a way to authorize activities other that fishing that result in the death of fish. Globally, most compensation activities involve creating or restoring habitats, which is often done in “like-for-like” schemes, where compensation attempts to replace areas of a given habitat lost to development by an equal or greater amount of the same habitat (McKenney and Kiesecker 2010; Quétier and Lavorel 2011). When “like-for-like” offsetting is not feasible, a different approach called “out-of-kind” offsetting is employed, where impacts to one population or location are compensated for by improving conditions for a different population, or by easing pressures from a different threat on the targeted population (Bull et al. 2013; Tallis et al. 2015). For example, fish mortality due to entrainment by water intake structures of a nuclear plant were proposed to be compensated by removing a dam 50 km inland from the facility (Barnthouse et al. 2019), and effects of restoration actions for compensating fishing mortality of American lobsters (French McCay et al. 2003) or seabird bycatch (Pascoe et al. 2011) were assessed using population models. The added challenge of “out-of-kind” offsetting is that losses and gains are often measured in different units or in different locations, as was the case when mortality of Golden eagles *(Aquila chrysaetos)* from wind energy development was offset by reducing lead poisoning from ammunition throughout Wyoming state (Cochrane et al. 2015). However, the potential for “out-of-kind” habitat compensation to offset anthropogenic mortality, by equating ongoing or chronic mortality to the habitat compensation required to appropriately mitigate population-level impacts, has not been previously evaluated.

Coho and other species of Pacific salmon *(Oncorhynchus* sp.) are ecologically, culturally, and economically important species in the PNW (Gende et al. 2002; Naiman et al. 2002), but have declined dramatically in recent decades (Slaney et al. 1996; Gustafson et al. 2007; Crozier et al. 2019). The main causes of decline include poor marine survival, commercial and recreational harvest, climate change, competition from hatchery fish, and the degradation of freshwater habitats needed for spawning and rearing (Kareiva et al. 2000; Chittenden et al. 2010). Degradation of freshwater habitats include the alteration of in-stream and adjacent terrestrial habitats from forestry practices (Scrivener and Brownlee 1989; Hartman et al. 1996; Tschaplinski and Pike 2017), the creation of barriers to migration, and the entrainment of fish in water intake structures of small and large hydropower dams (e.g. in the Columbia River system, Levin and Tolimieri 2001). As a consequence, an estimated $1 billion (US) a year has been spent on restoration of streams and rivers within the continental US since 1990 (Bernhardt et al. 2005). Here we use a matrix population model of coho salmon *(Oncorhynchus kisutch)* in the Pacific Northwest (PNW) of North America to evaluate the efficacy of “out-of-kind” offsetting and estimate the amount of compensation habitat required to achieve “No Net Loss” of productivity for salmon populations affected by anthropogenic development activities that cause mortality. Juvenile coho salmon rely on small streams as freshwater nursery habitat and off-channel habitats, like sloughs, side-channels, beaver ponds, or temporary to permanent floodplains, are often a target for restoration or compensation actions (Morley et al. 2005; Roni et al. 2006). Therefore, we evaluated how much compensation habitat (in m^2^), in the form of constructed off-channel habitat, is required to offset mortality on coho salmon and maintain overall population size and productivity. Further, we assessed how chronic anthropogenic mortality to three coho life history stages (eggs, parr, smolt/adult), individually and in combination, influenced overall population dynamics, and thus, the effectiveness of habitat compensation. Using threatened populations of coho salmon as an important case-study to develop a quantitative framework for balancing development losses and mitigation gains also serves the important call for research into addressing the uncertainty in offset analysis (Bull et al. 2013).

## Methods

### Scenarios

To evaluate the amount of habitat compensation required to maintain productivity for coho populations impacted by anthropogenic mortality, we modelled the impacts of chronic mortality and the offsetting potential of constructed compensation habitat for three life stages of coho salmon: egg, parr, and smolt/adult. We also evaluated how habitat compensation requirements change when mortality occurs at multiple life stages simultaneously. We focused on constructed side-channels because of their importance for rearing and overwintering coho juveniles (Morley et al. 2005; Rosenfeld et al. 2008), and because they are commonly used to limit or reverse the decline of anadromous salmon in the PNW (Nickelson and Lawson 1998; Ogston et al. 2015).

We expected that the value of constructed compensation habitats to the overall population dynamics of coho salmon would depend on which life history stage(s) experienced additional mortality. We created scenarios that varied the chronic mortality imposed on each life stage as well as our assumptions about the productivity of smolts from compensation habitats, and used a 3-stage deterministic matrix population model with density-dependent survival in the fry to parr transition to assess impacts to population size. The first set of scenarios (Egg scenarios) imposed mortality on eggs prior to density-dependent survival. Egg scenarios encompass a wide range of activities known to cause anthropogenic mortality to the egg stage, including increased sediment deposition or scour of spawning areas following deforestation (Scrivener and Brownlee 1989), thermal extremes (Tang et al. 1987), and dewatering of spawning areas below dams (Becker and Neitzel 1985, Casas-Mulet et al. 2005). In Parr scenarios, we represented chronic anthropogenic mortality caused by activities such as dam or powerplant entrainment (Muir et al. 2001; Barnthouse 2013), flow fluctuations due to hydropower generation (Nagrodski et al. 2012; Gibeau et al. 2017) and degradation in water quality from forestry practices or urbanization (Hartman et al. 1996; Chittenden et al. 2010). In Smolt/Adult scenarios, we imposed chronic mortality to represent impacts including smolt entrainment in, or passage over, dams (Levin and Tolimieri 2001; Schreck et al. 2006), or adult mortality during upstream migration due to warmer temperatures downstream of dams or powerplants (Baisez et al. 2011). In both the Parr and the Smolt/Adult scenarios, we assumed individuals experienced chronic anthropogenic mortality after the period of density-dependent survival (i.e., fry stage).

Precise estimates of mortality by classes of anthropogenic disturbance are rare, and relating stage-specific vital rates (e.g. survival) to degraded habitat conditions difficult (Hayes et al. 2009). Consequently, we simulated a broad range (2-20% per year) of chronic annual mortality to represent disturbances causing relatively small to large annual mortality (e.g. juvenile entrainment in spillways or turbines of dams, Muir et al. 2001). We applied annual mortality rates to egg, parr, and smolt/adult stages, both separately and in combination, to evaluate the potential population-level consequences of chronic anthropogenic impacts, and the scope for compensation habitats to ameliorate those effects.

### Deterministic matrix model

We created a deterministic 3-stage matrix model with a one-year time step to represent a typical 3-year coho salmon life-cycle (Nickelson and Lawson 1998; Bradford et al. 2000), and ran each simulation for 45 years to compare the final population size under baseline conditions to those for scenarios varying life stages affected, rates of chronic mortality, and sizes and productivity of compensation habitat. In our deterministic model, salmon spawn in late fall, while eggs incubate over winter, and fry emerge in the spring (Table 1). Fry quickly transform into juveniles called parr, which typically remain in freshwaters until their second spring, before migrating to the ocean as smolt. They finish their transformation to adulthood in the ocean, where they spend another 18 months before returning to freshwaters to spawn (Groot et al. 1995). The first stage (*F_13_*) in our model thus includes adult fecundity (*P_fem_*F_eggs_*), egg (*φ_em_*), and fry (*f_(φspr)_*) survival, while the second stage (*a_21_*) includes the early months of adult survival (*φ_oceY2_*) in the ocean. The third stage (*a_32_*) is made up of adult ocean survival as well as migration upstream into spawning habitats (*φ_oceY3_*).

**Table 1.**
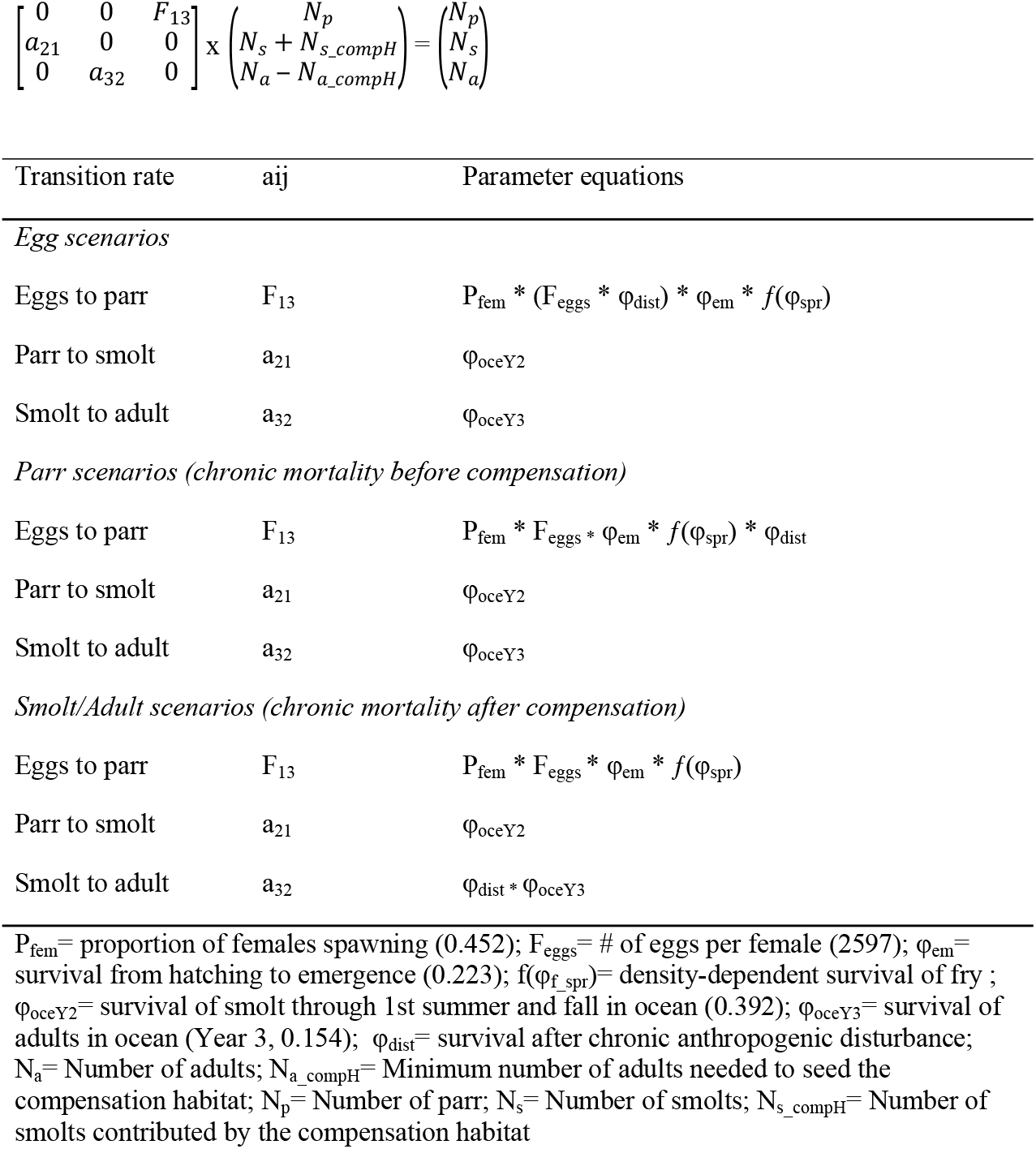
Deterministic matrix model and scenarios per life stage.

| Transition rate | a<sub>ij</sub> | Parameter equations |
| --- | --- | --- |
| *Egg scenarios* |  |  |
| Eggs to parr | F<sub>13</sub> | P<sub>fem</sub> * (F<sub>eggs</sub> * φ<sub>dist</sub>) * φ<sub>em</sub> * f(φ<sub>spr</sub>) |
| Parr to smolt | a<sub>21</sub> | φ<sub>oceY2</sub> |
| Smolt to adult | a<sub>32</sub> | φ<sub>oceY3</sub> |
| *Parr scenarios (chronic mortality before compensation)* |  |  |
| Eggs to parr | F<sub>13</sub> | P<sub>fem</sub> * F<sub>eggs</sub> * φ<sub>em</sub> * f(φ<sub>spr</sub>) * φ<sub>dist</sub> |
| Parr to smolt | a<sub>21</sub> | φ<sub>oceY2</sub> |
| Smolt to adult | a<sub>32</sub> | φ<sub>oceY3</sub> |
| *Smolt/Adult scenarios (chronic mortality after compensation)* |  |  |
| Eggs to parr | F<sub>13</sub> | P<sub>fem</sub> * F<sub>eggs</sub> * φ<sub>em</sub> * f(φ<sub>spr</sub>) |
| Parr to smolt | a<sub>21</sub> | φ<sub>oceY2</sub> |
| Smolt to adult | a<sub>32</sub> | φ<sub>dist</sub> * φ<sub>oceY3</sub> |

P<sub>fem</sub>= proportion of females spawning (0.452); F<sub>eggs</sub>= # of eggs per female (2597); φ<sub>em</sub>= survival from hatching to emergence (0.223); f(φ<sub>f\_spr</sub>)= density-dependent survival of fry ; φ<sub>oceY2</sub>= survival of smolt through 1st summer and fall in ocean (0.392); φ<sub>oceY3</sub>= survival of adults in ocean (Year 3, 0.154); φ<sub>dist</sub>= survival after chronic anthropogenic disturbance; N<sub>a</sub>= Number of adults; N<sub>a\_compH</sub>= Minimum number of adults needed to seed the compensation habitat; N<sub>p</sub>= Number of parr; N<sub>s</sub>= Number of smolts; N<sub>s\_compH</sub>= Number of smolts contributed by the compensation habitat

We assumed survival from fry to smolt was density-dependent with a bottleneck occurring immediately after fry emergence in the spring, since territorial fry compete for limited resources (Chapman 1962; Ward and Slaney 1993; Nislow et al. 2004). The density-dependent relationship was modelled with a Beverton-Holt function (Beverton and Holt 1957):

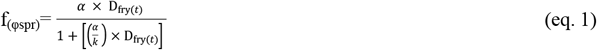

Where *α* is the number of smolts per fry at the origin, D_fry*(t)*_ is the number of fry at emergence (i.e. *P_fem_ * F_eggs_* * *φ_em_*), and *k* is the carrying capacity for smolts in the stream (smolts per km). We modeled freshwater density dependent survival after fry emergence by assuming *α* = 0.5 which is the average from 10 coho populations in the PNW (Bradford 1995; Korman and Tompkins 2014). We fixed the carrying capacity for smolts (*k*) at 15,318, corresponding to the mean *k* from field estimates (i.e. 1702 smolts/km, Bradford 1995; Korman and Tompkins 2014), multiplied by 9 km, the average length of the spawning or rearing reaches of 10 streams.

We ran the deterministic model in the absence of added anthropogenic mortality and density dependence to calculate the stable age distribution and the final population sizes under baseline conditions (808 adults, 13,379 parr, and 5249 smolts). We used the stable age distribution as the starting population vector for all subsequent simulations.

We assumed compensation side channels were fully functional immediately after construction, produced smolts at the maximum capacity each year, and the capacity to produce smolts remained constant over the length of the simulations. Spawning adults were first allocated to constructed side-channels until fully seeded (*N_a_compH_*, varied with the size of compensation simulated in each scenario), after which any remaining spawning adults were allocated to the main channel. We computed *N_a_compH_* by using average vital rates,

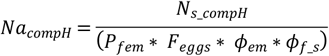

where *N_s_compH_* is the number of smolts contributed by the compensation habitat (varied with the size of side-channels in each scenario), *P_fem_* is the proportion of females returning to spawn (mean = 0.452), *F_eggs_* is the number of eggs produced per female (mean = 2597), *φ_em_* is the survival rate from eggs to fry emergence (mean = 0.223), and *φ_f_s_* is the average survival rate from fry to smolts in the absence of density dependence (0.075).

We calculated how many smolts would need to be produced from constructed compensation habitats in order to maintain population productivity (i.e. achieve offsetting equivalency), defined as population abundance returning to baseline (pre-impact) levels within 45 years. To do so, we ran simulations for each chronic mortality rate and impacted life stage over a range of compensation habitat sizes constructed for coho salmon in the PNW (Figure 1, *n* = 27 sites, Morley et al. 2005; Roni et al. 2006; Rosenfeld et al. 2008), and evaluated if offsetting equivalency was achieved in 45 years. In Parr scenarios, we assume that the compensation habitats were constructed downstream of the source of mortality so that smolts produced in compensation habitats were not exposed to the anthropogenic mortality. In comparison, in Smolt/Adult scenarios, chronic anthropogenic mortality impacted smolts or adults from both the compensation habitat and main population. Finally, we assessed the additional compensation requirements if chronic mortality occurred during both parr and smolt/adults stages. To do so, we ran simulations with *φ_dist_* applied to parr in the second year of the life cycle (*a21*), and to smolt/adults in the third year (*a32*).

**Figure 1.**
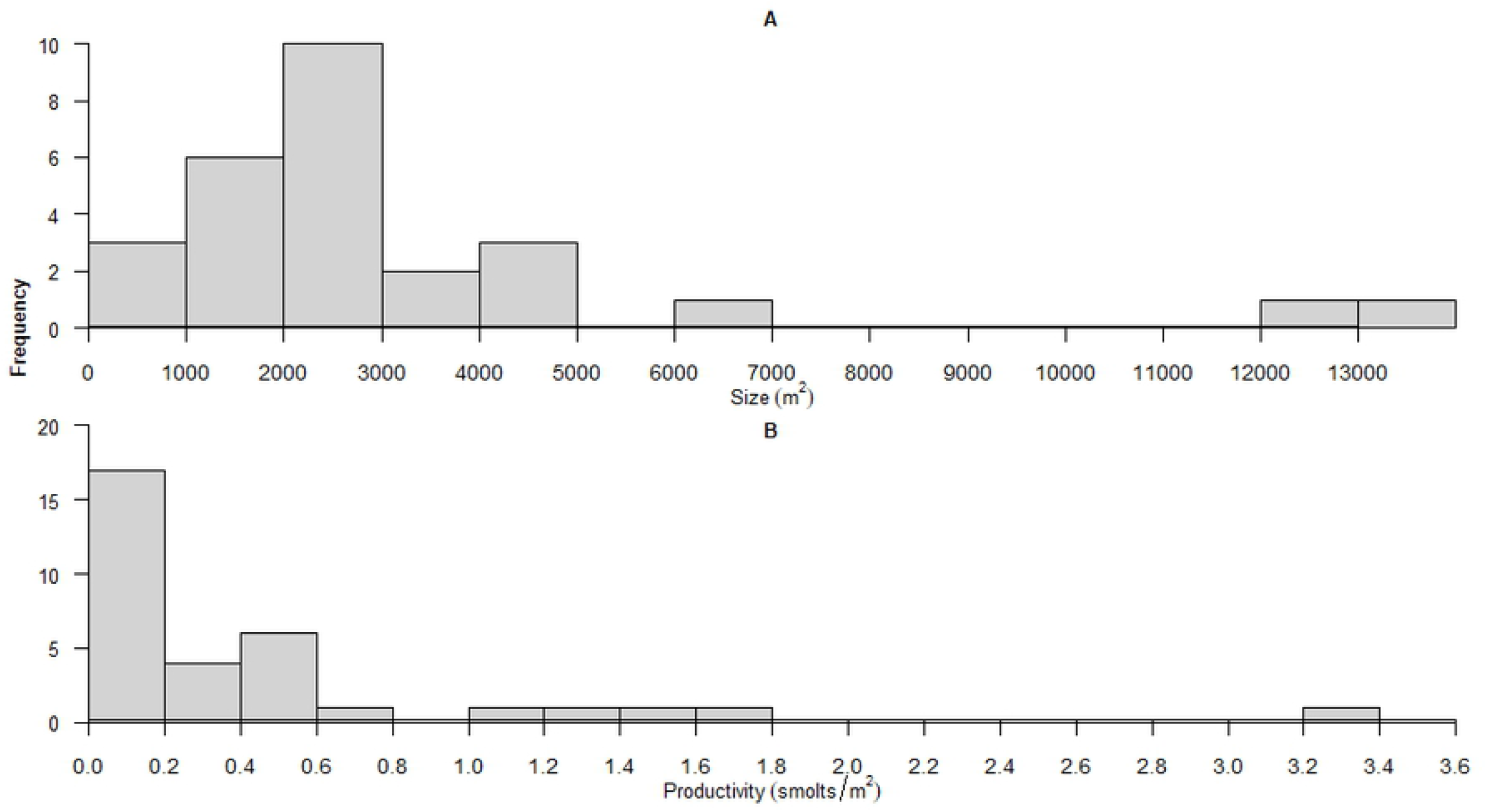
Distribution of (a) the size (m^2^) of 27 side-channel habitats built in the PNW (from Morley et al. 2005; Roni et al. 2006; Rosenfeld et al. 2008), and (b) the productivity of 33 side-channel habitats (# of smolts per m^2^) built in the PNW (from Rosenfeld et al. 2008; Roni et al. 2010).

### Compensation channels

Finally, our simulations also explored how assumptions about the productive capacity (i.e. quality, expressed as # of smolts produced per m^2^) of compensation habitats influenced the size of habitats needed to achieve offsetting equivalency. In the baseline scenario we assumed a mean smolt production in side-channels of 0.47 smolts per m^2^, corresponding to the mean production estimated from 33 constructed side-channels in the PNW (Rosenfeld et al. 2008; Roni et al. 2010; Figure 1.b). We also relaxed these assumptions and used the 25^th^ (0.1 smolt / m^2^), 50^th^ (0.18 smolts / m^2^), and 75^th^ (0.54 smolts /m^2^) percentiles of smolt production as additional scenarios.

All analyses were performed in program R (version 3.5.1, R Core Team 2013).

### Elasticity analyses

We performed a simulation-based elasticity analysis to assess how proportional changes in population growth rate changed relative to proportional changes in the individual vital rates included in the deterministic matrix model without density dependence (Morris and Doak 2002). We created 10,000 matrices with vital rates drawn at random from uniform distributions between the upper and lower 95% confidence interval for each vital rate, and calculated the deterministic growth rate (λ) (Wisdom et al. 2000). Density-dependent fry survival (*f*(_φspr_) was replaced by deterministic fry survival in year 1 (φ_fryY1_) and year 2 (φ_fryY2_), that averaged 7.5%, the mean fry-to-smolt survival across the 10 creeks used to derive the Beverton-Holt density-dependent function (citation). We used a general linear model with Gaussian distribution to decompose the variation in lambda, in which the standardized regression coefficients associated with each vital rate in the model estimate the proportional contribution of each vital rate to the variation observed in lambda (i.e. elasticity, Wisdom et al. 2000).

## Results

The population-level effect of chronic anthropogenic mortality depended on the life stage that experienced the mortality relative to the stage when density-dependent mortality occurred. When we modeled egg mortality without compensation, the impact on final population sizes was modest, with no more than a 3% reduction from the baseline population size. In comparison, when chronic mortality was imposed on parr or smolts/adults, after the period of freshwater density dependence, the final population size declined as chronic mortality increased (Figure 3).

**Figure 2.**
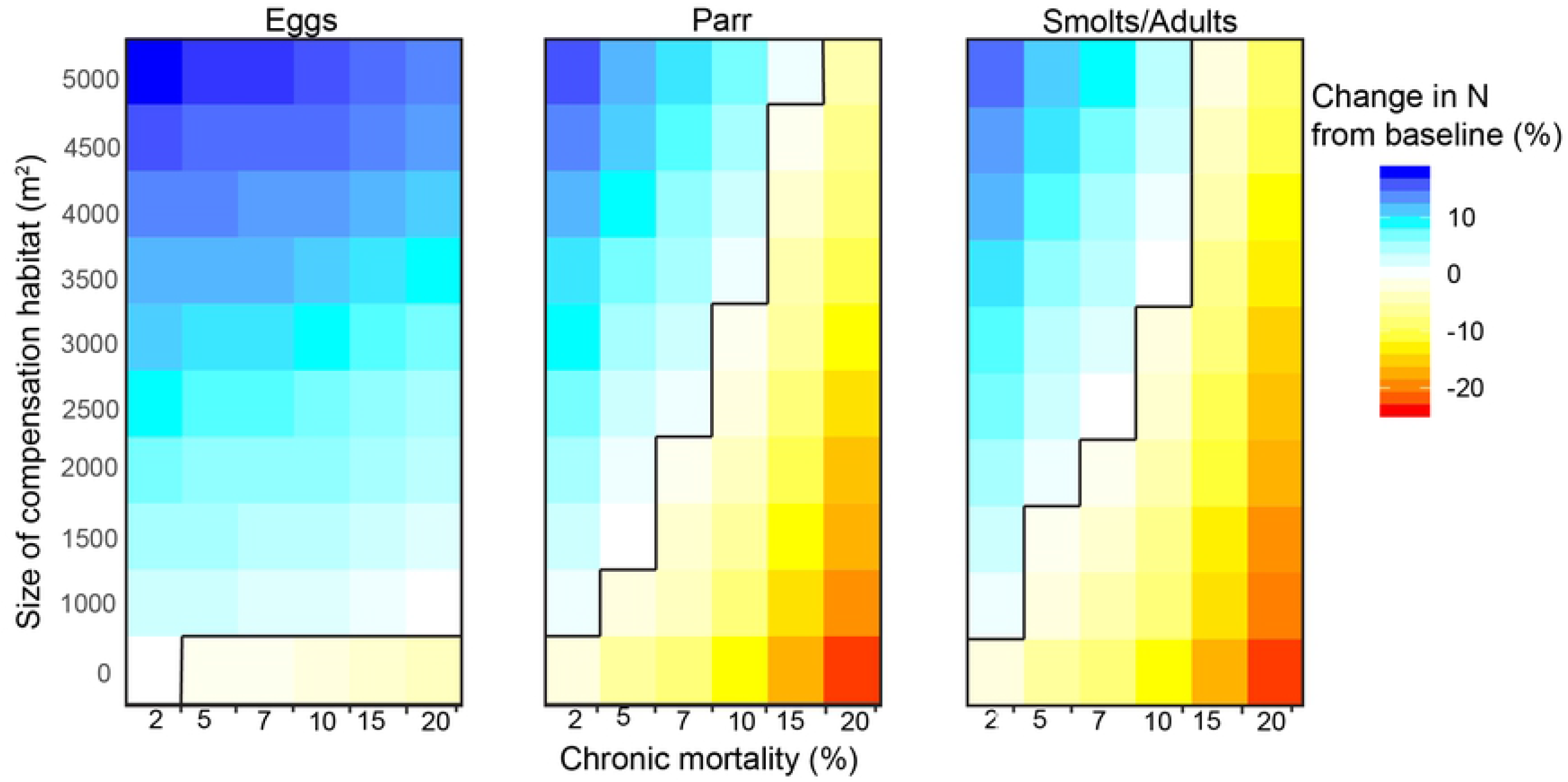
Change in final number of adults for impacted populations compared to baseline abundances (%, colors) over a range of sizes of constructed side-channels (y-axis), annual chronic anthropogenic mortality (x-axis), and life stage (egg, parr, smolt/adult) affected by the chronic mortality (panels), assuming mean productivity in constructed side-channels. 0 (white cells) indicates no change in final abundances of impacted populations compared to baseline, while positive values (cool shades) mean compensation increased the final number of adults in impacted populations and negative values (warm shades) mean final size of impacted populations decreased despite the amount of compensation habitat added. The black lines indicate offsetting equivalency for the combination of compensation habitat sizes and chronic mortality.

**Figure 3.**
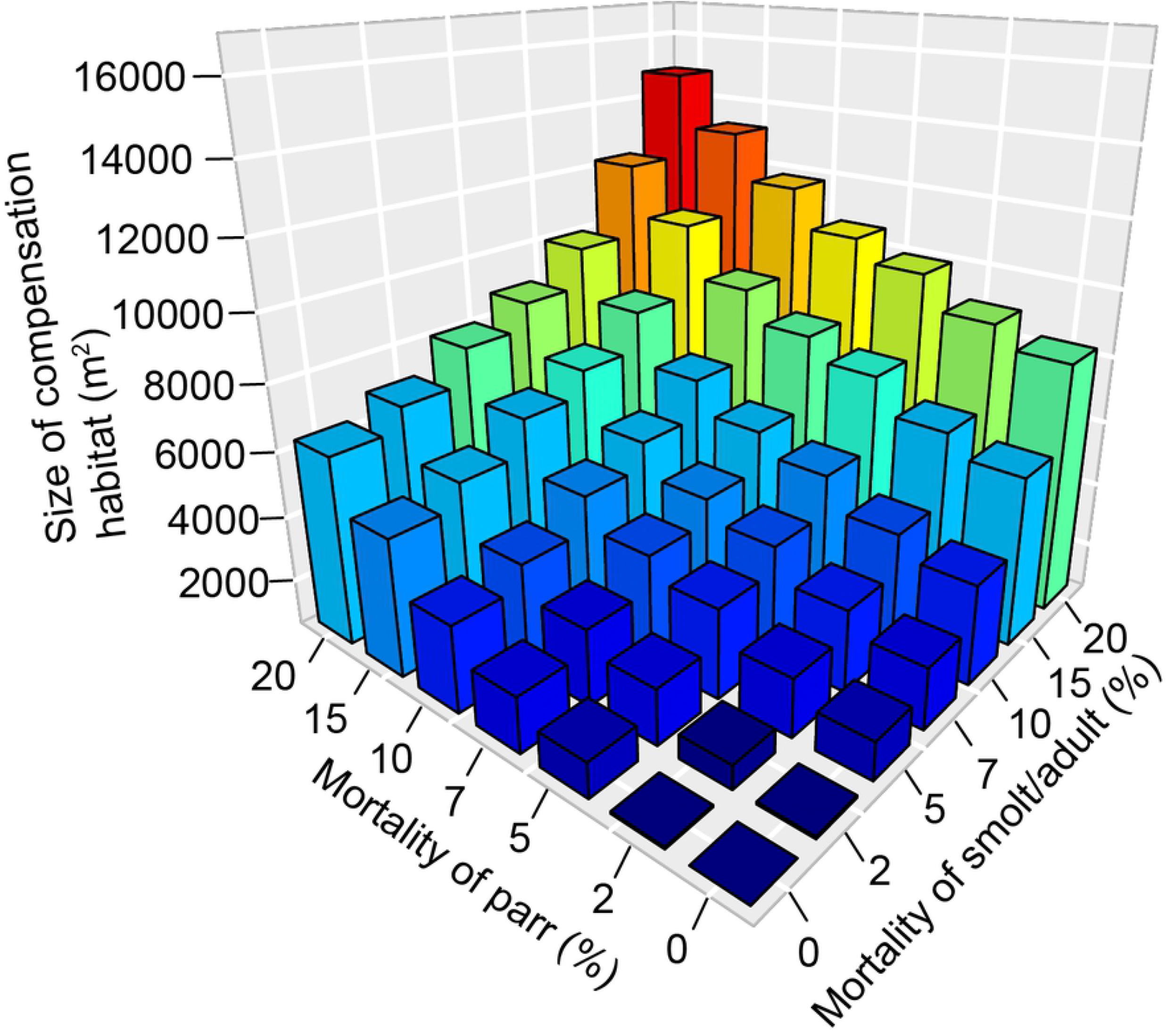
Range in size of compensation habitat (m^2^) (z-axis) required to effectively offset annual chronic mortality ranging from 0 to 20% when chronic mortality is applied cumulatively to both parr (x-axis) and smolt/adult (y-axis) life stages. We assumed mean productivity in constructed side-channels (0.47 smolts per m^2^).

**Figure 4.**
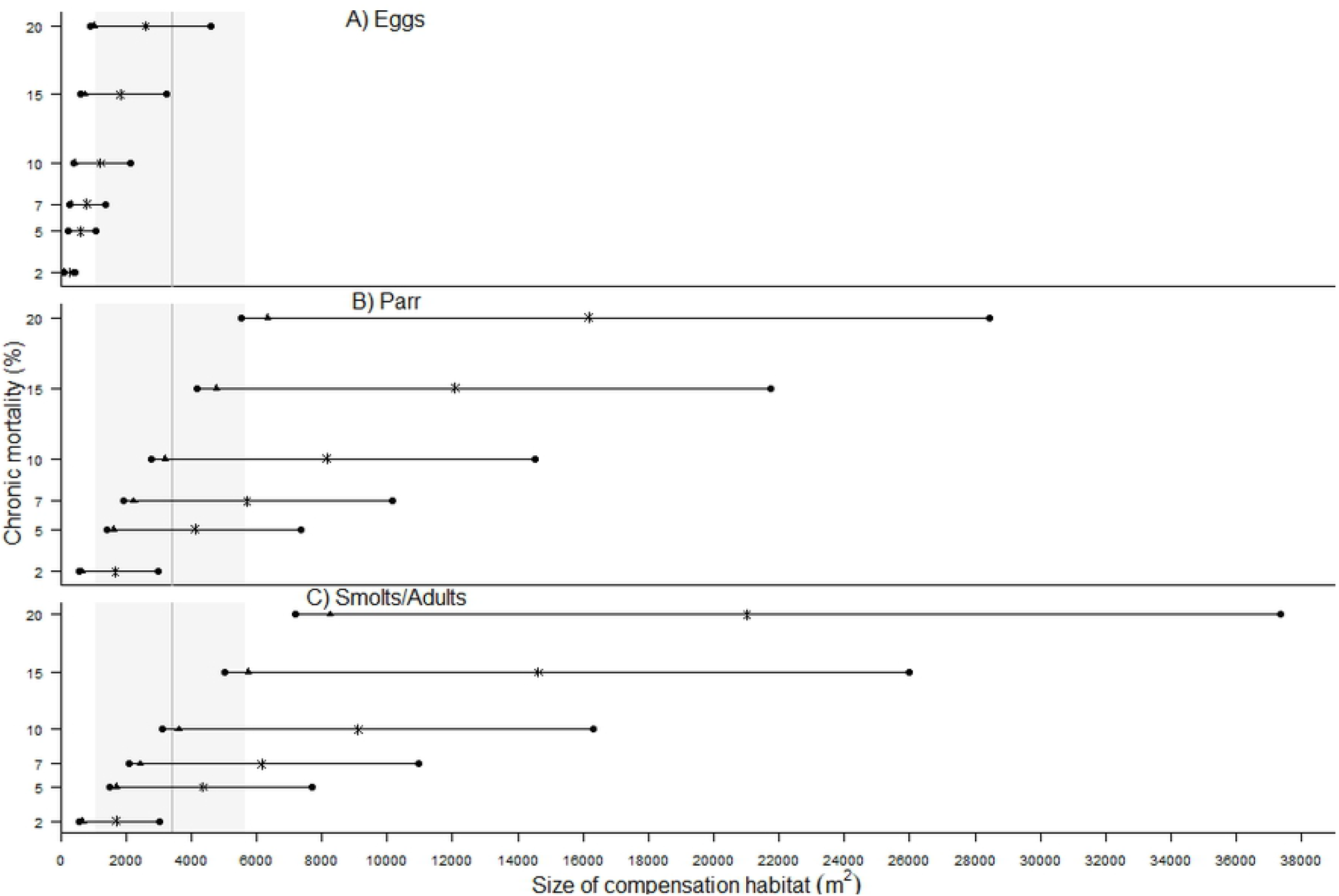
Range in sizes of compensation habitat (m^2^) required to effectively offset annual chronic mortality ranging from 2 to 20% affecting, a) Egg, b) Parr, and c) Smolt/Adult life stages, assuming varying productivity (# of smolts produced per m^2^) for the constructed side-channels. Horizontal black lines indicate the 25^th^ to 75^th^ percentiles of productivity of compensation habitat, black triangles represent mean number of smolts contributed, and stars represent median number of smolts contributed by compensation habitat (based on data from 33 sites, from Rosenfeld et al. 2008; Roni et al. 2010). The grey box represents the mean (vertical line) and 10^th^ to 90^th^ percentiles (shaded) of sizes of compensation habitats built in the PNW (n= 27 sites, data from Morley et al. 2005; from Roni et al. 2006; Rosenfeld et al. 2008).

Simulated populations that experienced chronic anthropogenic mortality benefitted from the addition of smolts from constructed side-channels, regardless of the life stage affected (Figure 3). We found that final population sizes increased linearly with the size of constructed side-channels until, and beyond, achieving offsetting equivalency (i.e. reaching adult abundances similar to baseline levels). However, we found that the importance of side-channel compensation habitat varied depending on the timing of mortality in the coho life cycle. For example, a relatively small constructed side-channel was sufficient to achieve offsetting equivalency when mortality impacted the egg stage, but the size of side-channels required to offset mortality occurring on the later life stages was much greater. For example, a small side-channel of approximately 1000 m^2^ (i.e., smaller than the 10^th^ percentile of side-channels built in the PNW) could compensate for up to 20% chronic annual mortality to the egg stage, but only 2% chronic annual mortality to parr or smolts/adults (Figure 3). Generally, more compensation habitat was needed to achieve offsetting equivalency when smolts/adults were affected compared to parr, especially if the intensity of chronic mortality was greater than 5% annually (Figure 3).

When chronic mortality was applied to both parr and smolt/adult life stages, the size of compensation habitat needed to reach offsetting equivalency increased with the magnitude of chronic mortality (Figure 2. Change in final number of adults for impacted populations compared to baseline abundances (%, colors) over a range of sizes of constructed side-channels (y-axis), annual chronic anthropogenic mortality (x-axis), and life stage (egg, parr, smolt/adult) affected by the chronic mortality (panels), assuming mean productivity in constructed side-channels. 0 (white cells) indicates no change in final abundances of impacted populations compared to baseline, while positive values (cool shades) mean compensation increased the final number of adults in impacted populations and negative values (warm shades) mean final size of impacted populations decreased despite the amount of compensation habitat added. The black lines indicate offsetting equivalency for the combination of compensation habitat sizes and chronic mortality.

Figure). The cumulative, or combined, effect of extra mortality on parr and smolt/adult stages was largely additive, but became multiplicative when anthropogenic mortality to smolt/adults was greater than 7%, and mortality to parr above 10%. For example, when an additional 20% mortality was applied to both parr and smolt/adults stages annually, the size of constructed side-channels needed to achieve equivalency increased by 1.8 to 2.3 times (from 6,336 m^2^ or 8259 m^2^, respectively, to 14,867 m^2^) compared to when the same chronic mortality was applied to either the parr or smolt/adult stage separately.

### Productivity of compensation habitats

Our results suggest that assumptions regarding the productivity of compensation habitats (i.e., the number of smolts produced by the constructed side-channels) have a large effect on our conclusions regarding the size of compensation needed to achieve offsetting equivalency. Based on the 25^th^ percentile of productivity, the size of side-channels required to achieve offsetting equivalency was 4.5 times greater than if we assumed mean quality (Figure).

#### Elasticity analyses

Our simulation-based elasticity analyses assessed how population growth rates changed relative to proportional changes in vital rates and indicated that survival of fry to smolt (*φ_f_s_* elasticity of 0.15) and adult ocean survival (elasticity of 0.16) had the largest influence on deterministic population growth rates. These two rates were three times more important than fecundity (*Feggs*) and the survival of fry at emergence (elasticity of 0.051, 0.049, respectively), while the proportion of females *(Pfem)* had very little influence on population growth (elasticity of 0.0007).

## Discussion

We used a matrix population model to compare how mortality at one life stage can or cannot be offset by adding production of the same or a different life stage from compensation habitat, in an equivalency analysis using an “out-of-kind” scheme to meet requirement for No Net Loss of population productivity. Our models suggest that the average size of constructed side-channels typically built in the PNW could compensate for chronic mortality of up to 20% annually if it affected the egg stage, but only up to 7% if it affected parr, smolts, or adults. Averaged-size side-channels would not be sufficient if added annual mortality was greater than 10% for parr or smolt/adult stages, greater than 5% if both parr and smolt/adult stages were affected, or if the productivity of side channels is lower than the average values used in our baseline scenario. Constructed side-channels have a wide range of productivities as a result of differences in design and site-specific considerations (Rosenfeld et al. 2008; Roni et al. 2010). Thus, a more precautionary approach to building side-channels would assume less than ideal productivity in compensation habitats. If we assumed lower productivity, side-channels would need to be bigger than those required when we assumed mean productivity (*sensu* Rosenfeld et al. 2008; Roni et al. 2010). Additionally, our results suggest that compensation habitats are more effective at mitigating chronic mortality if they are built downstream of where disturbances occur (i.e., as in the parr scenarios), such that smolts they produce are not affected by chronic mortality experienced by the population in the main channel (which happens in the smolt/adult scenarios). Overall, if chronic mortality also affects the smolts contributed by the compensation habitats (e.g. in cases of smolts entrainment into downstream dams or increased ocean mortality), achieving offsetting equivalency will require more compensation habitat to be built in freshwater.

As our modelling of an “out-of-kind” compensation scheme highlights, designing effective side-channels for compensation is complex. Our results emphasize the importance of considering variation in productivity when deciding on the optimal size of compensation habitat needed. However, the size of constructed side-channels may also impact quality of the compensation habitat. For example, studies have noted a decline in smolt density with increasing side-channel sizes, suggesting that smaller side-channels may be more demographically efficient than large side-channels (Keeley et al. 1996; Rosenfeld et al. 2008). Building numerous, but smaller, side-channels may also be more technically and economically feasible than fewer large ones. In our models, we assumed that larger compensation habitats could be equally as productive as smaller habitats, but if productivity declines with increasing size or through time due to degradation, our results may underestimate the sizes needed to maintain population equivalency.

Adding to the complexity of designing constructed side-channels of adequate size and quality, the size and the number of side-channels are frequently chosen based on availability of land and costs of restoration rather than based on ecological bottlenecks or the potential for success (Rosenfeld et al. 2008). Such an opportunistic selection of compensation sites is stated as one reason why “No Net Loss” of productivity is not often achieved in practice (Quigley and Harper 2006). Our results suggest that intentionally siting side-channels downstream of where anthropogenic impacts occur would be more effective at compensating for chronic mortality.

However, downstream reaches of streams and rivers are often less stable geomorphologically (Naiman et al. 2000; Naiman et al. 2008), and the substantial investment required to build compensation habitats in low-elevation floodplains could be lost if, or when, natural large flow events occur. Other challenges to building effective side-channels habitats include the need for connectivity between the compensation habitats and main channel, which is crucial to ensure the success of mitigation (Anderson et al. 2014; Crozier et al. 2019). For example, schooling of juveniles and limited migration may lower the carrying capacity of constructed side-channels by limiting the number of fry that move in from spawning or rearing grounds (Walters et al. 2013). Salmon also need dynamic and diverse freshwater habitats to thrive, and compensation projects will be more effective if they consider natural evolutionary processes needed for the health and resilience of the species (Waples et al. 2009; Beechie et al. 2010; Booth et al. 2016), especially considering the added challenges posed by accelerating climate change. For example, improving temperature and flow regimes through riparian restoration and increased food availability may improve the tolerance of salmon populations to warmer water temperatures induced by climate change (Crozier et al. 2019). Our results support the conclusion that cumulative impacts from multiple sources of anthropogenic mortality on more than one life-stage can have compounding (i.e. multiplicative) effects on population dynamics, and increase the need for mitigation and compensation activities. As such, our simulations of only a single life stage experiencing chronic mortality are likely to be underestimates of the minimum sizes of constructed side-channels needed to maintain population sizes in the face of multiple, overlapping sources of anthropogenic mortality (e.g. Hartman et al. 1996).

At the population level, our results highlight that the potential for effective mitigation of added chronic mortality depends both on understanding the influence of density-dependent survival in affected populations, as well as on the unequal contributions of different life stages to the overall population dynamics. For example, natural density-dependent bottlenecks may compensate for some mortality caused by anthropogenic disturbances if it occurs prior to periods of negative density dependent survival (Hodgson et al. 2017), and influence the need for compensation habitat. Our results showing only modest population-level impacts of additional egg mortality illustrate the low elasticity of the egg stage to affect overall population dynamics, as well as the potential for density-dependent survival to compensate for some anthropogenic mortality. Moreover, our elasticity analysis highlights the large influence that variation in fry to smolt survival (elasticity of 0.15) has on population growth, separate from the issue of density dependence, further emphasizing the importance of survival at that stage for salmon conservation efforts. However, it is risky to rely on natural density dependence to compensate for chronic anthropogenic mortality as it remains very challenging to detect if, when and how strong density-dependent survival bottlenecks occur in freshwater for specific populations (Rose et al. 2001; Milner et al. 2003; Walters et al. 2013). Moreover, density-dependent survival may only compensate for additional mortality when population sizes are large (e.g. approaching carrying capacity, Goodwin et al. 2006), which may be uncommon for many anadromous salmon populations currently impacted by anthropogenic disturbances (Achord et al. 2003), unless such disturbances also lower freshwater carrying capacity (Greene and Beechie 2004). Other forms of density dependence may enhance the efficacy of compensation habitat in mitigating the effects of disturbances. For example, density-dependent migration, whereby individuals in areas with high densities move to areas with lower densities, allows the compensation habitat to be more effectively populated by encouraging more individuals to colonize it (Greene and Beechie 2004). Overall, given the difficulty in adequately assessing the presence, timing, and strength of density-dependent survival in freshwater, long-term studies of population dynamics of local populations appear crucial to adequately design effective compensation habitats. Finally, a substantial amount of uncertainty remains in the application of our modelling results due to the difficulty in estimating mortality rates for various sources of anthropogenic mortality. Quantifying mortality from human activities (e.g. forestry, entrainment in dam structures, harmful discharges etc.), along with a method such as ours for how stage-specific mortality translates into population-level impacts, is crucial information needed for calculating offsetting requirements.

## Conclusion

Our results suggest that compensation habitats built for coho salmon in freshwater have the potential to mitigate some chronic and ongoing mortality. However, our results also indicate that achieving offsetting equivalency may require creating a much larger amount of side-channel habitat than is typically constructed in the PNW (Rosenfeld et al. 2008; Roni et al. 2010), especially when we relaxed assumptions about ideal smolt productivity in constructed side-channels, or considered cumulative impacts to multiple life stages. Moreover, our results illustrate how life-cycle population models can be a powerful tool to examine the efficacy of restoration efforts targeted to different life stages for overall population dynamics. We show that matrix population models can be used to quantitatively estimate the uncertainty in “out-of-kind” offset analysis, though their utility can be limited by the availability of site-specific information on population productivity and the magnitude of additional mortality. Protecting and restoring threatened salmon populations in the PNW requires a critical evaluation of the efficacy of policies and practices of industry and regulators. Overall, the importance of side-channels for juvenile coho salmon and the large influence of fry-to-smolt survival rates on overall population dynamics suggested by our elasticity analysis highlight the value of building side-channels as compensation habitat for coho salmon, despite the complexity and challenges involved.

## Acknowledgements

We thank K. Wilson, D.A Greenberg, A. Cantin, R. Murray, and J.W. Moore for help and comments that greatly improved the manuscript. This work was funded by grants from NSERC Discovery Grant, the Gordon and Betty Moore, and Wilburforce Foundations to WJP.

